# Parallel transcriptome profiling of soybean cyst nematodes (SCN) during a three-way interaction among SCN, soybean aphids, and soybean

**DOI:** 10.64898/2026.11.09.765379

**Authors:** Bibek Adhikari, Surendra Neupane, Adam J. Varenhorst, Shyam Solanki, Madhav P. Nepal

## Abstract

The soybean cyst nematode (SCN), *Heterodera glycines* (Ichinoe) (Rhabditida: Heteroderidae) is arguably the greatest yield-limiting factor for soybean (*Glycine max* [L.] Merr.) production in the United States. Previous research found that the impact of SCN infestation on yield is exacerbated during concurrent infestations with soybean aphids (SBA), *Aphis glycines* (Matsurmura) (Hemiptera: Aphididae). Although recent research has characterized the host root transcriptomic response during these infestations little is known about the corresponding in planta gene expression dynamics of either pest. This study examines the transcriptomic expression of SCN in soybean cultivars’ roots under single or combined SBA infestations at two different time points: 5 and 30 days post-infestation (DPI). The SCN transcriptomics for SCN-susceptible (S) PI 518671 (Williams 82) cultivar reacted equivalently to both SCN and SCN + SBA treatments. However, transcriptomics in SCN-resistant (R) MN1806CN (*H. glycines* HG type 0/race 3 resistant) demonstrated an elevated significant expression pattern under the SCN + SBA combined treatment. During the early infestation period (i.e., sampled at 5 DPI), both S and R cultivars reacted similarly, irrespective of treatment conditions. The functional enrichment analysis of 1829 differentially expressed genes/transcripts showed that during the early co-infestation period (5 DPI), SCN genes were significant and mapped towards resistance breakage, whereas in the late stage (30 DPI), the gene flux oriented towards nematode dormancy. Together, these results suggest that SCN activity is amplified when combined with SBA, offering foundational data for molecular breeding to produce SCN resistant soybean cultivars.

## 1. Introduction

Soybeans (*Glycine max* [L.] Merr.), rich in high-quality protein and oil, are among the four major global commodity crops, alongside wheat, corn, and rice [1,2]. The United States is the second-largest producer of soybeans, accounting for a 27% share of global production following Brazil (42%) [3]. However, diseases and pests account for production losses, ca. 4.8% of potential soybean production was lost to diseases in 2025 in the US alone [4]. The soybean cyst nematode (SCN), *Heterodera glycines* (Ichinohe) (Rhabditida: Heteroderidae), is the most economically important below-ground pest infesting soybean roots [5]. Similarly, soybean aphid (SBA), *Aphis glycines* (Matsumura) (Hemiptera: Aphididae), is an invasive phloem feeder that is an economically important aboveground soybean pest [6]. In the US, the annual cost of SCN infestation on soybean yield exceeds $1.5 billion [7,8], meanwhile between $949 million and $1.623 billion is attributed to SBA [9].

Noticing the impact of pests, SBA- and SCN-resistant soybean genotypes are widely classified and commercially available for cultivation practices [10,11]. However, the cultivar resistant to one pest might be susceptible to the other and combined infestation often intensifies the yield loss [12]. Co-infesting SCN-susceptible (DK 28–52, IA 3018, IA 3041) and SCN-resistant (DK 27–52, AG 2821 V, IA 3028) soybean cultivars with SBA, fungus *Cadophora gregata*, and SCN, McCarville et al. [13] observed reproductive advantages to SCN, increasing population by ca. five times. The aboveground feeding by SBA in SCN-resistant (Dekalb 27–52, PI 88788 derived) and SCN-susceptible (Kenwood 94) cultivars improved the susceptibility to SCN [14]. However, preceding research by Neupane et. al [15,16] highlighted SCN reproductive advantage in the susceptible (Williams 82, PI 518671) cultivar and an unchanged effect in the resistant (MN1806CN, resistant to *H. glycines* HG type 0/race 3) cultivar under combined infestation. Collectively, these studies indicate that multi-pest co-infestations can alter host physiological responses and promote pest fecundity in a genotype-dependent manner. Consequently, elucidating the molecular mechanisms governing plant-defense signaling and pest virulence under concurrent biotic stress is essential for developing durable, multi-stress-tolerant cultivars.

The molecular expression of SCN genes inside the host has rarely been evaluated. Here, we studied the differential gene expression of SCN inoculated into susceptible and resistant soybean cultivars under single or SBA co-infested conditions, shifting from our previous insights from a host-to-pathogen perspective. Together, these findings offer necessary baseline data to elucidate the molecular pathogenicity of SCN, promoting potential molecular breeding initiatives to produce SCN-resistant soybean cultivars.

## 2. Materials and methods

### 2.1. Plant material and pest infestations

SCN-susceptible: PI 518671 (Williams 82) and SCN-resistant: MN1806CN (*H. glycines* HG type 0/race 3 resistant) were two soybean cultivars used in the study. The SCN and SBA populations obtained from two sources were reared in susceptible soybean hosts before greenhouse experimentation. HG type 0/race 3, SCN population, avirulence with respect to major SCN resistance sources and defined by <10% reproduction on indicator lines was maintained and populated on SCN-susceptible soybean. While SBA biotype-1 colonies obtained from The Ohio State University were maintained on a susceptible soybean line, LD12-15838R.

Details on randomization, experimental design, and greenhouse experimentation have been well described previously [15]. Here we focus on the soybean genotypes receiving the following treatments: no pest as control, SCNs only, and SCNs + SBAs. The 5 and 30 days post-infestation (DPI) root tissues were harvested, snap-frozen in liquid nitrogen and stored at −80°C before total RNA extraction. PureLink RNA Mini Kit (Invitrogen, Carlsbad, CA, USA) was used in total RNA extraction, treating with TURBO DNase (Invitrogen), subjected to 1% agarose gel electrophoresis assessment, and quantified using NanoDrop 2000 (Thermo Fisher Scientific, Waltham, MA, USA). The NEBNext Ultra II RNA Library Prep Kit (96 single index) was used in cDNA library preparation and sent to 100 bp single-end sequencing at the Iowa State University Sequencing Facility, IA, on an Illumina HiSeq 3000 platform.

### 2.2. RNA-seq preprocessing, mapping, quantification, and normalization

The raw sequence reads associated with BioProject PRJNA514200 (SRA study/Project ID: SRP178193) were obtained from NCBI Sequence Read Archive (SRA) using a custom script. The associated accession numbers under investigation are shown in Supplementary **Table 1**. FastQC (v0.11.9) [17] was used to assess raw read quality, while MultiQC (v1.13) [18] was employed for summarization. Btrim (v0.2.0) [19] was employed to remove low-quality (quality threshold >20; 5-bp sliding window) bases.

The SCN genome, transcript, and protein files were obtained from WormBase Parasite [20], accessed from https://parasite.wormbase.org/Heterodera_glycines_prjna381081/Info/Index (Last accessed on 25 August 2026). High-quality reads obtained after quality assessments were thus mapped against the coding sequences of SCN (heterodera_glycines.PRJNA381081.WBPS19.mRNA_transcripts.fa) using Salmon tool v2.6.0 [21], accessed through the Bioconda channel [22]. The txdbmaker [23] and AnnotationDbi version 1.74.0 [24] packages from the Bioconductor v3.23 [25] channel were employed in R 4.6.1 [26] using RStudio 2026.08.2 to map Salmon-quantified transcripts against the *H. glycines*’ gene (heterodera_glycines.PRJNA381081.WBPS19.annotations.gff3).

### 2.3. Differential gene expression analyses

Benjamini–Hochberg procedure with raw *p*-values adjusted for multiple testing was employed to control the false-discovery rate (FDR) [27]. In RStudio using base R, Bioconductor’s tximport [28] package was used for importing Salmon’s transcript-abundance reads and the DESeq2 v1.52.0 [29] package was employed to identify the differentially expressed genes (DEGs). The reads were filtered with a threshold > 10 counts, setting an absolute fold change (|FC|) ≥ 2 and FDR ≤ 0.01 for DEGs. The pheatmap version 1.0.13 [30] was employed to create heatmaps of differentially expressed genes, while ggplot2 version 4.0.3 [31] was used for principal component analysis visualization. Other R packages used in analyses include grid version 4.6.1, gridExtra version 2.3.1 [32], tidyr version 1.3.2 [33], dplyr version 1.2.1 [34], RColorBrewer version 1.1-3 [35], and VennDiagram [36].

### 2.4. Functional enrichment analyses

Protein sequences from “heterodera_glycines.PRJNA381081.WBPS19.proteins.fa” file, as observed in the significant DEGs .csv file, were extracted using Biostrings [37] package version 2.80.1. The extracted protein fasta files were subjected to eggNOG-mapper v3.0.0-beta6 [38] using orthology assignments from the eggNOG 7 database [39] for functional annotation. Homology searches were carried out with DIAMOND [40] in sensitive mode with iterative search enabled (ortholog e-value ≤ 0.001). Functional terms—Gene Ontology (GO) and Kyoto Encyclopedia of Genes and Genomes (KEGG) orthologs (KO)—were transferred from the identified orthologs within the automatically adjusted taxonomic scope (--tax_scope auto).

The GO terms obtained from eggNOG-mapper were employed to dotplot visualization utilizing readxl version 1.6.0 [41], dplyr, tidyr, stringr version 1.6.0 [42], AnnotationDbi, GO.db version 3.23.1 [43], and ggplot2 libraries. The KO hits were validated by searching KO terms in the KEGG database [44] through https://www.kegg.jp/kegg/ko.html (last accessed on 4 September, 2026).

## 3. Results

### 3.1. RNA-seq dataset quality and overall structure of the transcriptome signal

After adapter trimming and quality control, more than 99% of reads (%GC: ∼43–45) were retained, but the mapping rate was exceptionally low. While no mapping to the control was as expected, the low mapping was likely contributed to by higher host transcript expression or suppression of the pathogen by the resistant soybean genotype. **Table 1** provides the summary statistics of transcript reads. Initially, the reads were mapped against 23,933 *H. glycines* transcripts; later, they were converted to 22,465 genes for DEG analysis. A fully balanced 2 (genotype) x 3 (treatment) x 2 (day) design, considering biological replicates per group, for 35 samples, was employed for DEG analysis. After filtering out genes with a count threshold below ten, 12,951 reads were retained for differential expression testing. Principal Component Analysis (PCA), based on variance-stabilizing transformed (VST) top 2000 genes, indicated treatment as a dominant driver of expression variation (PC1 = 87.50%), while genotype contributed fractional variation (PC2 = 1.90%). This highlights that the treatment, rather than genotype or sampling day, was the dominant axis of transcriptional variation in this dataset (**Figure 1**).

**Figure 1.**
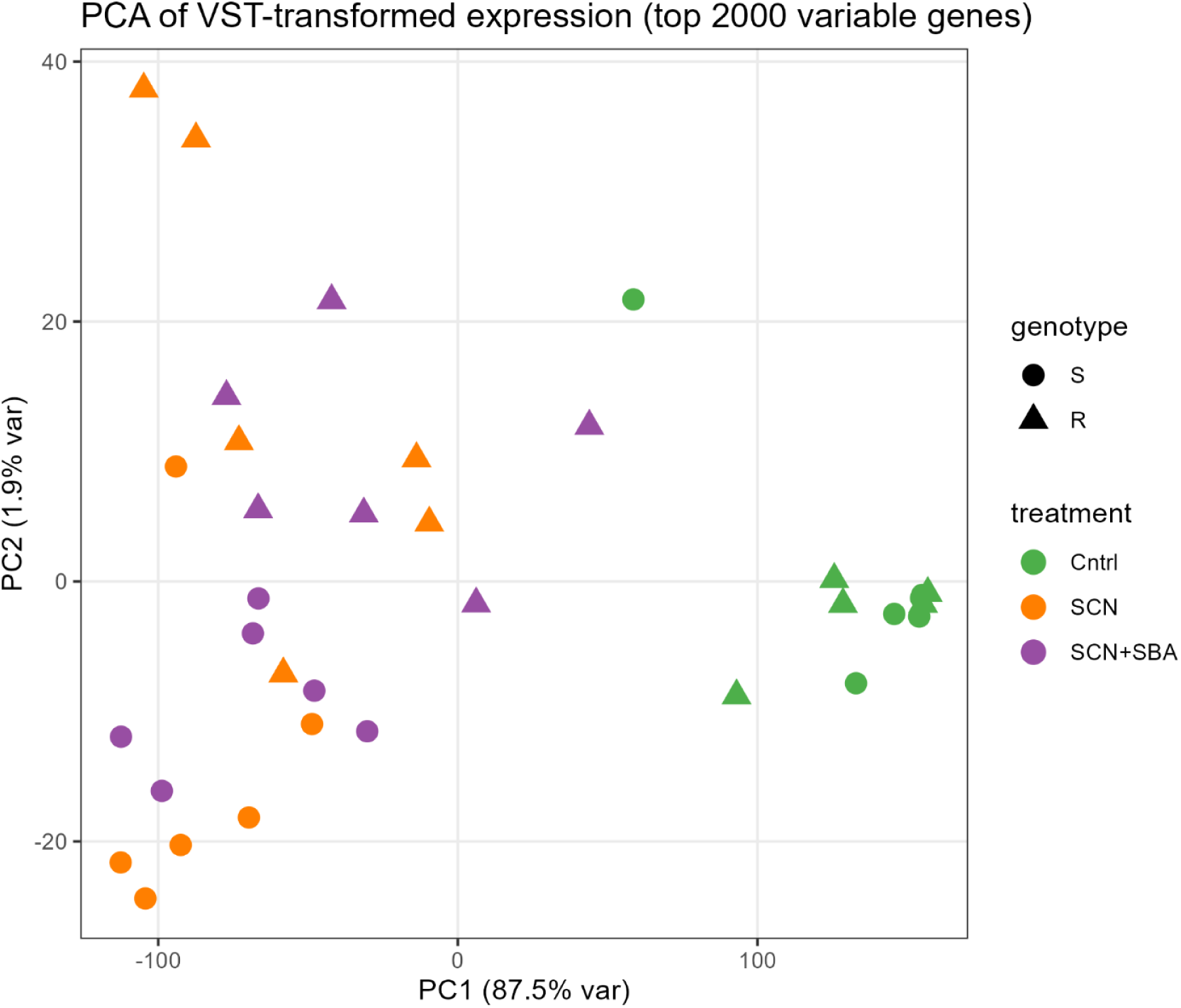
Principal Component Analysis (PCA), based on the top 2,000 variance-stabilizing transformed (VST) genes, plot showing separation of genotypes based on expressed SCN genes along with treatments. Each point represents an individual sample, where different shapes represent soybean genotypes, with colors corresponding to treatment. PC1 explains 87.50% of the observed variance while PC2 explains 1.90%.

**Table 1.** Summary table with significant DEGs expressed under different treatment conditions compared to genotype and time point.

| **Treatment** | **Genotype** | **DPI** | **Significant DEGs** | **Comparison** |
|---|---|---|---|---|
| SCN | S | 5 vs 30 | 0 | S_SCN_5vs30_DPI |
| SCN vs SCN+SBA | S | 5 | 0 | S_SCNvsSCN+SBA_5_DPI |
| SCN vs SCN+SBA | S | 30 | 0 | S_SCNvsSCN+SBA_30_DPI |
| SCN+SBA | S | 5 vs 30 | 2 | S_SCN+SBA_5vs30_DPI |
| SCN | R | 5 vs 30 | 870 | R_SCN_5vs30_DPI |
| SCN vs SCN+SBA | R | 5 | 0 | R_SCNvsSCN+SBA_5_DPI |
| SCN vs SCN+SBA | R | 30 | 112 | R_SCNvsSCN+SBA_30_DPI |
| SCN+SBA | R | 5 vs 30 | 1127 | R_SCN+SBA_5vs30_DPI |
| SCN | R vs S | 5 | 3 | RvsS_SCN_5DPI |
| SCN+SBA | R vs S | 5 | 63 | RvsS_SCN+SBA_5DPI |
| SCN | R vs S | 30 | 1 | RvsS_SCN_30DPI |
| SCN+SBA | R vs S | 30 | 105 | RvsS_SCN+SBA_30DPI |

### 3.2. Genotype shaping the transcriptome’s significant expression

To test whether resistant (R) and susceptible (S) genotypes differ transcriptionally independent of treatments, we compared R versus S separately at five and 30 DPI, modeling treatment as a covariate (design: ∼treatment + genotype). Using a Benjamin-Hochberg adjusted p-value (padj) <0.05 and an absolute log2 fold-change >2, we identified 1,829 significantly regulated genes across the genotypes (**Figure 2**). While the controls were evaluated for non-DEGs, the heatmap in **Figure 2** shows significant DEGs across genotype comparisons. The only two significant DEGs in the susceptible genotype treated with SCN + SBA, compared between five and 30 DPI, include: Hetgly15078 (log2FC = -5.3853, padj = 0.0041) and Hetgly15081 (log2FC = -4.9787, padj = 0.000347).

**Figure 2.**
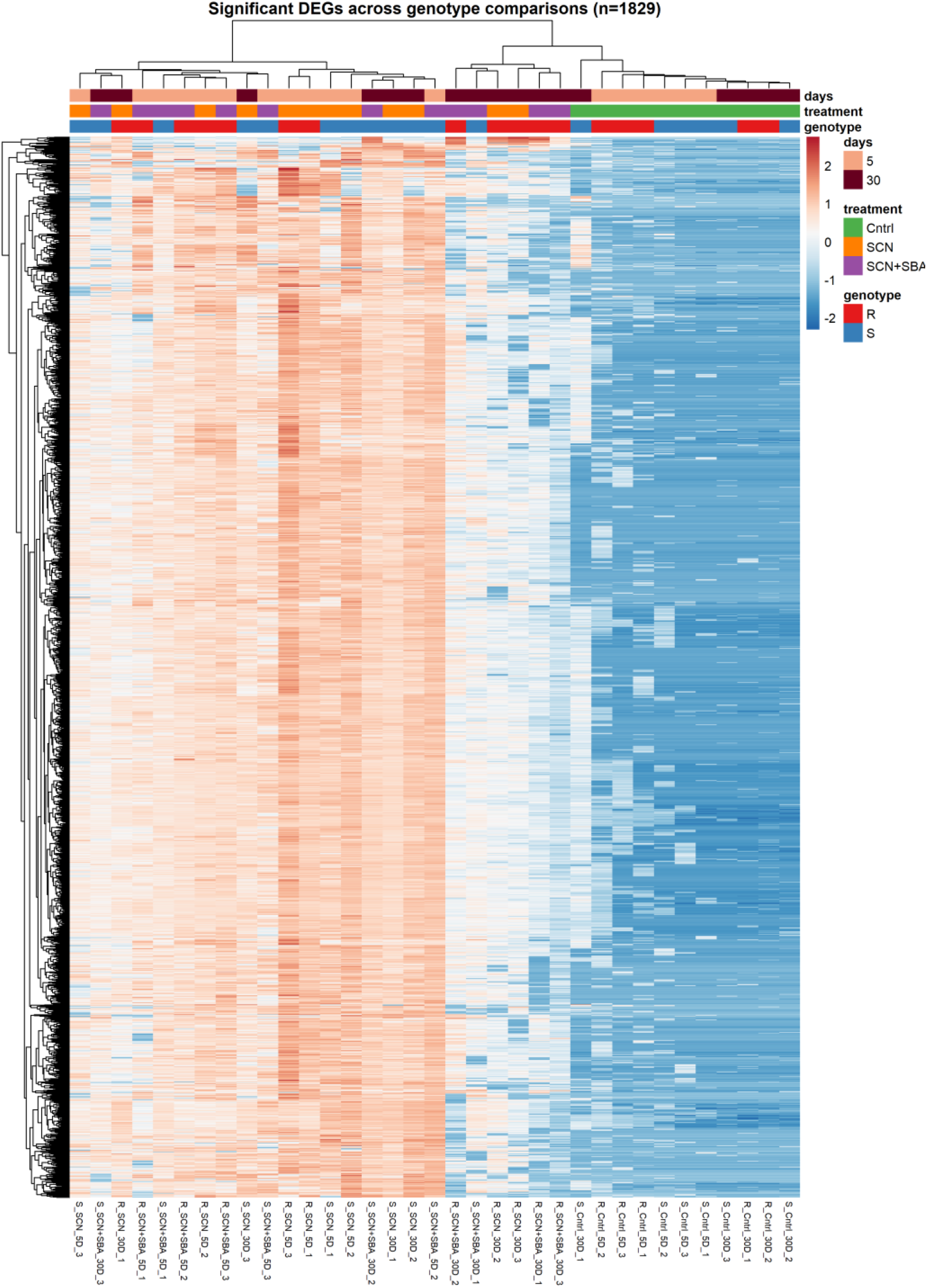
Heatmap showing the significant DEGs (n=1829) across soybean genotypes and treatments. The branches at the top highlight the sample clusters, while those on the left denote gene clusters. The denser red and light blue bars represent over- and under-expressed genes. Different color codes on the top panel represent days, treatments, and genotypes, as shown in the legend. The sample name is at the bottom, where S = susceptible, R = resistant, Cntrl = control, SCN = soybean cyst nematode, SCN + SBA = coinfection of SCN with soybean aphid, 5D and 30D = 5^th^ and 30^th^ days post-infection, and the last number as replicates.

For the five vs 30 DPI comparison of resistant cultivar infested with SCN, Hetgly03580 (log2FC = 8.6258, padj = 3.42E-06), Hetgly21679 (log2FC = 7.1445, padj = 0.0002), Hetgly21220 (log2FC = 7.1298, padj = 0.0078), Hetgly21462 (log2FC = 6.9996, padj = 0.0136), Hetgly07143 (log2FC = 6.4822, padj = 0.0016), Hetgly21034 (log2FC = 6.4372, padj = 0.0120), Hetgly10380 (log2FC = 6.2394, padj = 0.0009), Hetgly01940 (log2FC = 5.7190, padj = 0.0093), Hetgly16253 (log2FC = 5.6992, padj = 0.0369), and Hetgly13787 (log2FC = 5.36138, padj = 0.0131) are the top 10 genes with log2FC > 5; while Hetgly15081 (log2FC = -5.3402, padj = 8.19E-06), Hetgly20925 (log2FC = -5.8703, padj = 0.0272), Hetgly19367 (log2FC = -6.2303, padj = 0.0242), Hetgly12699 (log2FC = -7.9574, padj = 0.0106), and Hetgly05084 (log2FC = -9.5933, padj = 0.0214) are the top five genes with log2FC < -5. For the 5 vs 30 DPI comparison of resistant cultivar infested with SCN + SBA, top three upregulated (log2FC > 10) genes include: Hetgly05515 (log2FC = 46.33412, padj = 7.18E-30), Hetgly00187 (log2FC = 45.84539, padj = 2.87E-85), and Hetgly02176 (log2FC = 10.70806, padj = 0.000171), while top three downregulated (log2FC < -5) genes include Hetgly07468 (log2FC = -5.20255, padj = 0.0207), Hetgly15078 (log2FC = -5.83146, padj = 0.0001), and Hetgly12699 (log2FC = -7.97719, padj = 0.0075).

### 3.3. The resistant genotype responded robustly

We next examined treatment effects within each genotype separately (expression: ∼ genotype x treatment x time), testing eight pairwise contrasts: SCN versus SCN + SBA at each time point, and the change from five to 30 DPI within each treatment for both genotypes (**Table 1**). In the susceptible genotype, only one contrast yielded two significant DEGs (SCN + SBA, five vs. 30 DPI), indicating negligible transcriptional variation to SCN or SCN + SBA treatment, or to time, under the applied significance threshold. In marked contrast, the resistant genotype showed a clear, time-dependent response: 870 DEGs distinguished SCN treatment at five DPI from 30, 112 DEGs distinguished SCN from SCN+SBA treatment at 30 DPI, and 1,127 DEGs distinguished SCN + SBA treatment at five DPI from 30; the largest DEG set among the eight within-genotype contrasts. The volcano plots and Venn diagram in **Figure 3** illustrate significantly regulated DEGs for the resistant genotype. Additionally, when comparing DEGs between resistant and susceptible genotypes for SCN infestation, we observed only five and one gene at 5 and 30 DPI, respectively. Whereas for SCN + SBA coinfestation, 63 and 105 genes (5 vs. 30 DPI) were significantly expressed. This asymmetry suggests that the resistant genotype mounts a substantially more dynamic transcriptional response to nematode challenge and to the addition of SNA than the susceptible genotype. Ninety-one genes were shared between SCN vs SCN + SBA (30 DPI) and SCN + SBA (5 vs 30 DPI) sets, while only four genes were shared between the SCN (5 vs 30 DPI) and SCN vs SCN + SBA (30 DPI) sets, 180 between SCN (5 vs 30 DPI) and SCN + SBA (5 vs 30 DPI), and three genes were common to all three. This pattern indicates that the SCN + SBA time-course response and the 30 DPI SCN vs SCN + SBA response shared a substantial subset of genes. The three commonly significantly expressed genes for the resistant genotype across treatments and time points included Hetgly01075, Hetgly07954, and Hetgly08253, respectively.

**Figure 3.**
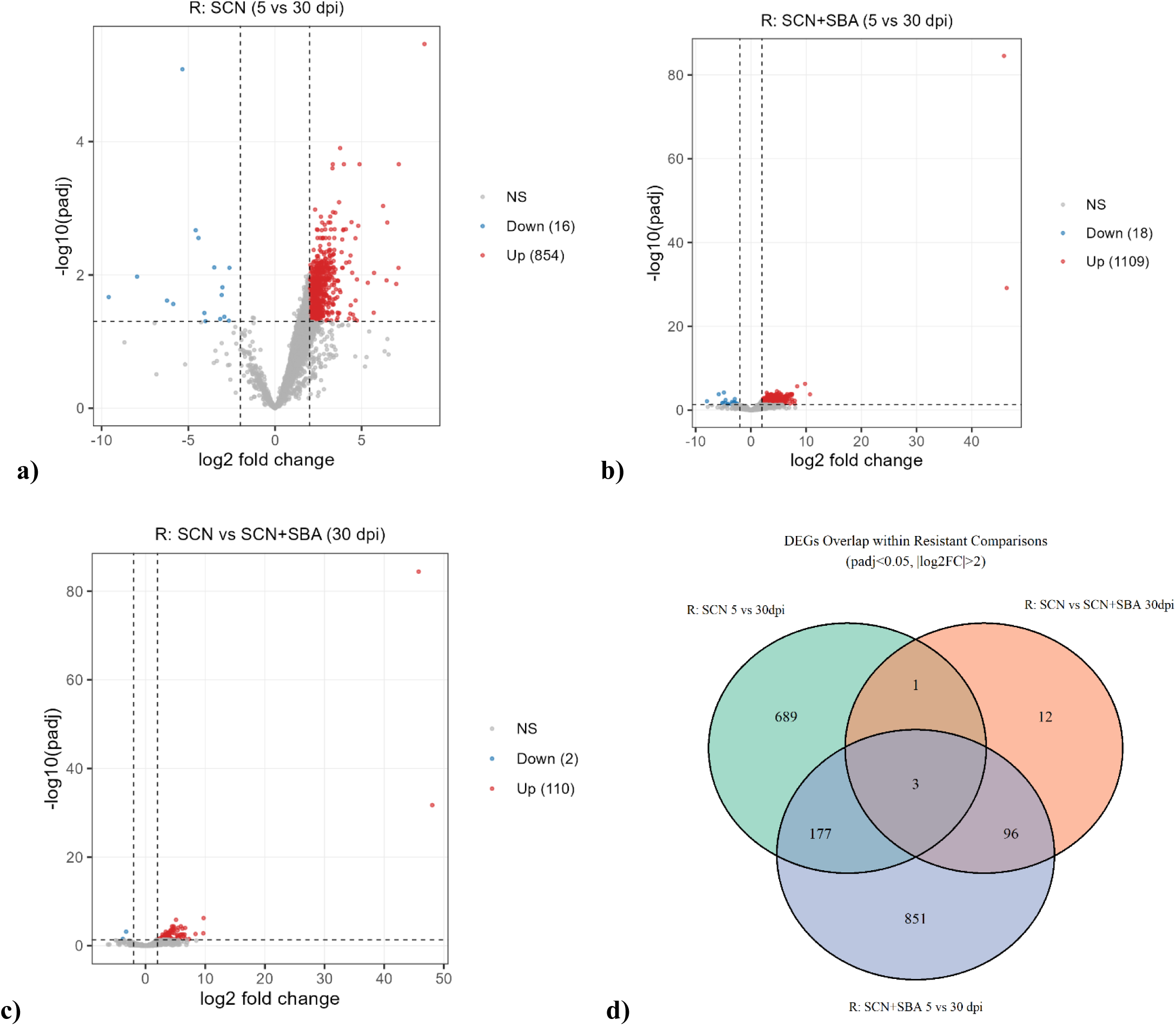
Differentially expressed genes for the resistant genotype. Volcano plots: (a) SCN (5 vs 30 DPI), (b) SCN + SBA (5 vs 30 DPI), and (c) vs SCN + SBA (30 DPI). Venn diagram (d) illustrating overlapping genes for the same conditions.

### 3.4. Solo and combined effect on resistant genotype

We compared SCN-only and combined SCN + SBA effects on the genotypes over time. For SCN-only treatment, very few genes were observed to be significantly expressed (three vs one at five and 30 DPI). Whereas when combined with SBA, the number of significantly expressed genes increased from 63 to 105 (5 vs 30 DPI). **Figure 4** illustrates the top 100 DEGs for SCN and SCN + SBA effects over time in the resistant genotype. When comparing resistant and susceptible cultivars for significantly expressed genes treated with SCN: Hetgly03580 (log2FC = 10.7841 and padj = 0.0155), Hetgly13322 (log2FC = 15.5218 and padj = 0.0174), and Hetgly19525 (log2FC = 4.5264 and padj = 0.0456) were only the three genes at five DPI; while Hetgly19250 (log2FC = 32.0865 and padj = 5.01E-05) was the only gene at 30 DPI.

**Figure 4.**
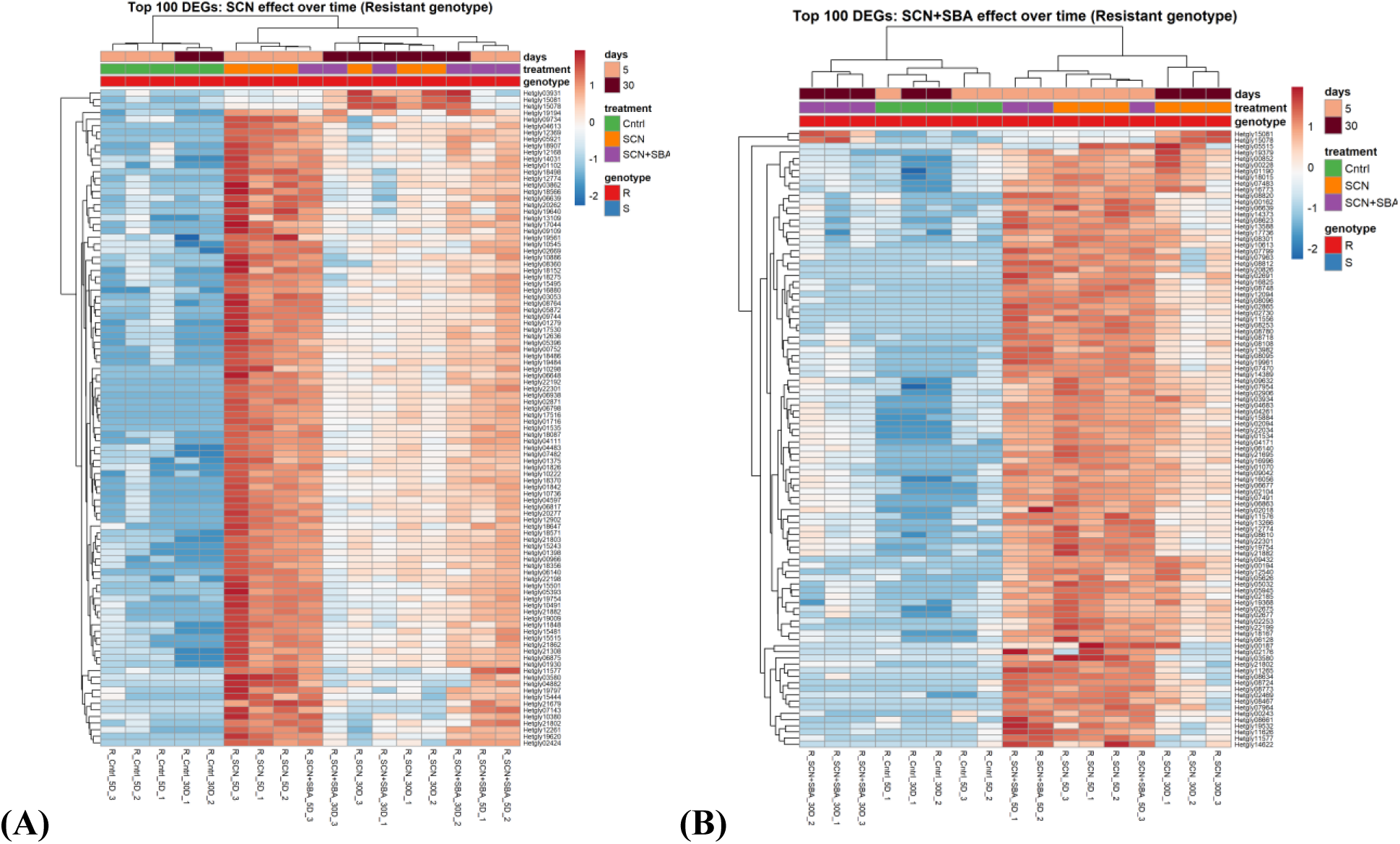
Heatmaps highlighting the significant DEGs across resistant genotypes as factor of treatments over time: **(A)** SCN and **(B)** combined SCN + SBA effects. The branches on the top show sample clusters, while those on the left denote gene clusters. The denser red and light blue colors represent over- and under-expressed genes. Different color codes in the top panel represent days, treatment, and genotype, as shown in the legend. The sample name is at the bottom, where R = resistant, Cntrl = control, SCN = soybean cyst nematode, SCN + SBA = coinfection of SCN with soybean aphid, 5D and 30D = 5^th^ and 30^th^ days post-infection, and the last number as replicates.

In contrast, for SCN + SBA treatment, out of 63 DEGs at 5 DPI, Hetgly13322 (log2FC = 6.4036, padj = 0.0182), Hetgly08764 (log2FC = 5.1727, padj = 0.0180), and Hetgly04988 (log2FC = 5.0185, padj = 0.0057) were top three upregulated genes (log2FC > 5); while Hetgly20245 (log2FC = -2.0140, padj = 0.0499), Hetgly02373 (log2FC = -3.3932, padj = 0.0147), Hetgly07978 (log2FC = -4.5715, padj = 1.13E-06), and Hetgly04944 (log2FC = -6.1386, padj = 0.0126) are the only downregulated genes. Similarly, at 30 DPI, out of 105 significant DEGs for SCN + SBA treatment, Hetgly19250 (log2FC = 14.8744, padj = 0.0006) and Hetgly19194 (log2FC = 2.6764, padj0.0029) were the only two upregulated genes; while Hetgly15984 (log2FC = -7.0213, padj = 0.0157), Hetgly08050 (log2FC = -7.0400, padj = 0.0002), Hetgly17077 (log2FC = -7.5243, padj = 0.0004), Hetgly02623 (log2FC = -7.8872, padj = 0.0003), Hetgly02845 (log2FC = -8.2139, padj = 0.0057), and Hetgly09432 (log2FC = -9.8793, padj = 2.40E-06) are top six downregulated genes (log2FC < -7). Hetgly02373 and Hetgly07978 were the common DEGs compared between resistant and susceptible cultivars treated with SCN + SBA (5 vs 30 DPI).

### 3.5. Functional association of DEGs

GO enrichment analysis of the proteins associated with significant DEGs in the resistant genotype returned 4,960, 4,669, and 601 GO terms, for SCN infestation at five vs 30DPI, SCN + SBA infestation at five vs 30dpi, and SCN vs SCN + SBA infestation at 30DPI treatment contrasts, corresponding to their predicted functions. The top eight GO terms associated with Biological Process, Cellular Component, and Molecular Function are visualized in **Figure 5A**-**C**. For SCN infestation at 5 vs 30 dpi and SCN + SBA infestation at five vs 30 DPI, the associated GO terms belonging to DEGs were similar. The top two GO associated with biological process include embryo development ending in birth or egg hatching (GO:0009792) and determination of adult lifespan (GO:0008340). The cellular compartment highlighted the majority of DEGs to nucleus (GO:0005634) and cytoplasm (GO:0005737) localized. Additionally, more than half of the DEGs highlighted protein binding (GO:0005515 and GO:0042802) molecular function. Even though the cellular compartment and molecular function for SCN vs SCN + SBA infestation at 30 DPI was similar to previous two contrasts, biological process included DEGs associated with collagen and cuticulin-based cuticle development (GO:0040002).

**Figure 5.**
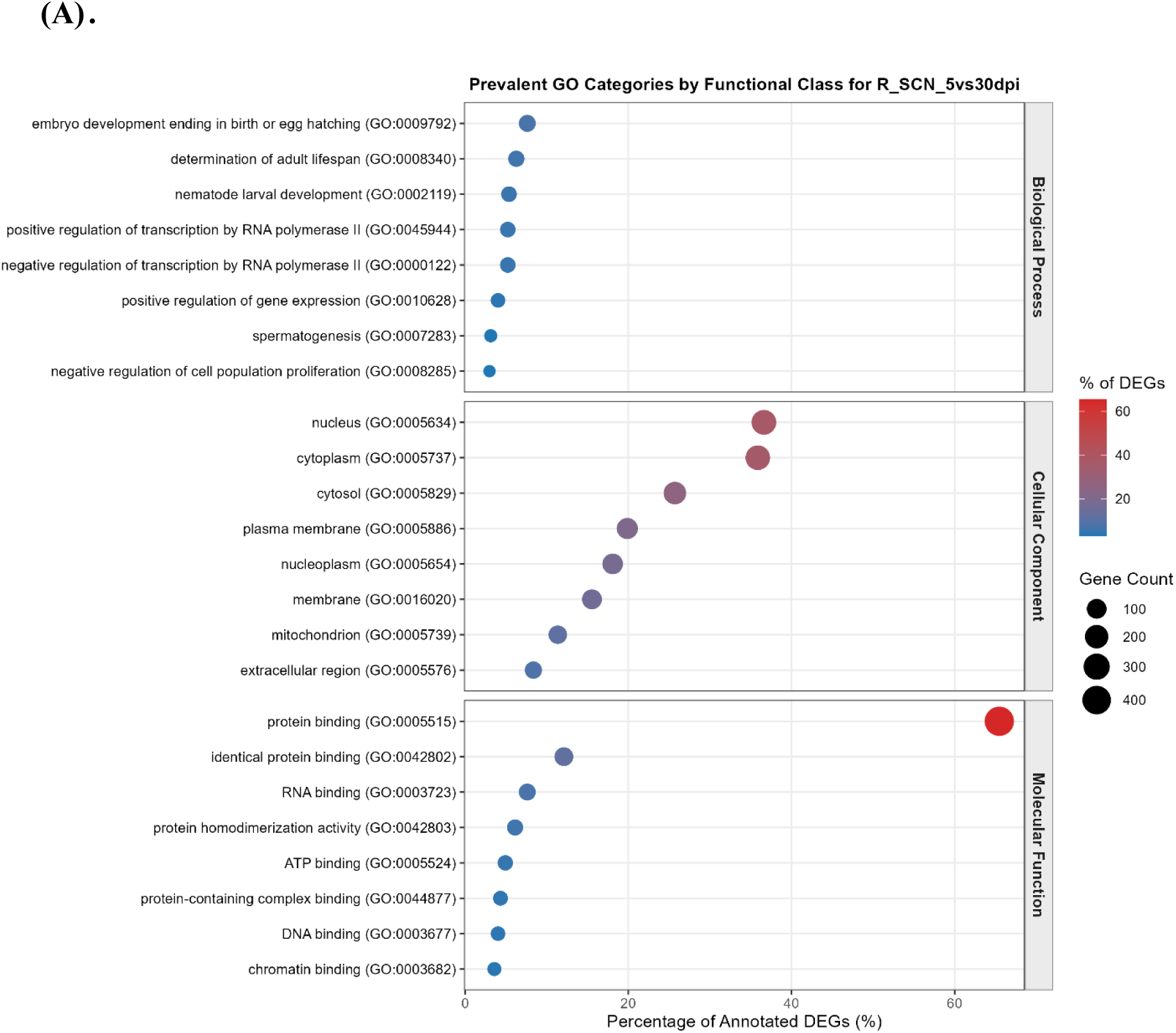
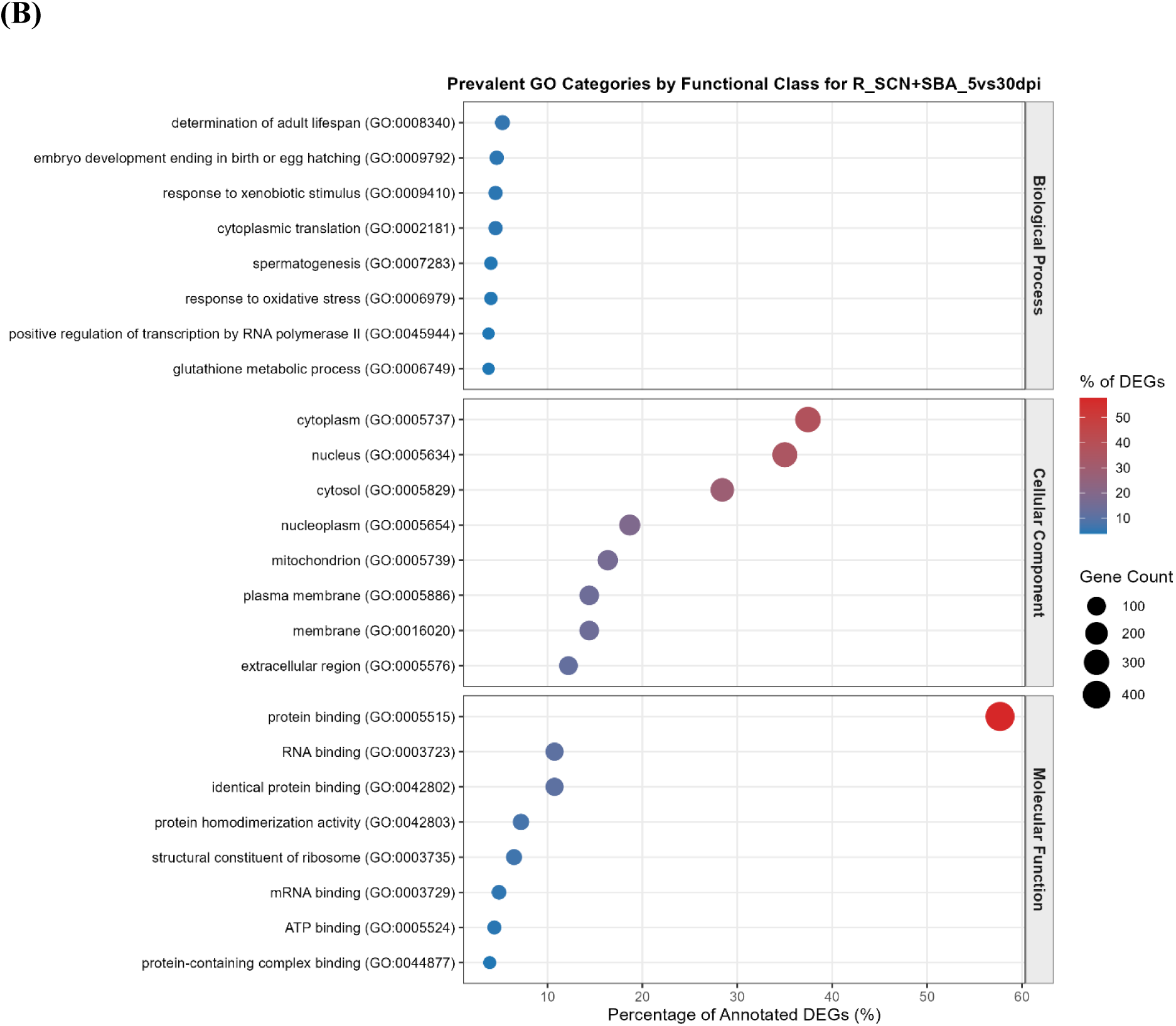
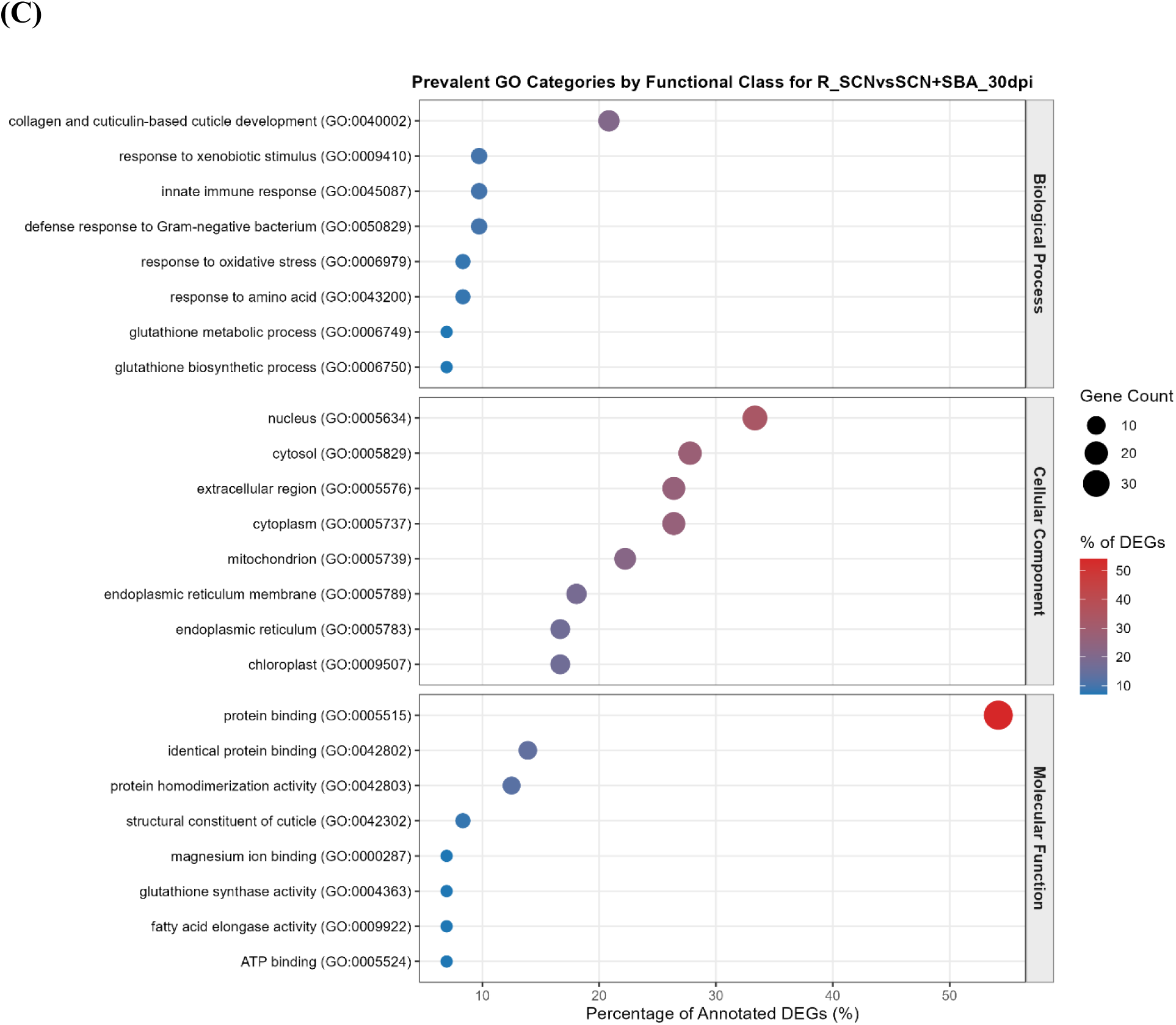
Gene Ontology (GO) annotations for DEGs in the resistant genotype. The dotplots illustrates top eight GOs associated with Biological Process, Cellular Component, and Molecular Function for **(A)** SCN infestation at five vs 30 DPI, **(B)** SCN + SBA infestation at five vs 30 DPI, and **(C)** SCN vs SCN + SBA infestation at 30 DPI. The x-axis represents percentage of annotated DEGs, while the y-axis represents the associated GOs. Different color bars represent % of DEGs and circle represents gene counts, as shown in the legend.

For SCN treatment, comparing five vs 30 DPI, the highly enriched KEGG Ortholog (KO) was K11513 (n=7) associated with centromere protein U, while transcript Hetgly21344.t1 (n = 15) was most repetitive. Similarly, when comparing SCN + SBA at 5 vs 30 DPI, K21456 (n = 22), associated with glutathione synthase, was highly enriched, while Hetgly07813.t1 (n =16) had the highest hits. Additionally, K08202 (n = 14), belonging to the MFS transporter [OCT family, solute carrier family 22 (organic cation transporter)], and Hetgly04034.t1 (n=8) were enriched KO and transcript, for SCN vs SCN + SBA at 30 DPI.

Similarly, GO analysis of the proteins associated with significant DEGs compared between resistant and susceptible genotypes resulted in 398 and 571 GO terms corresponding to their predicted functions for SCN + SBA infestation at 5dpi and SCN + SBA infestation at 30 DPI, respectively. The top eight GO terms associated with Biological Process, Cellular Component, and Molecular Function are visualized in **Figure 6A**-**B**. For five DPI contrast biological process include response to jasmonic (GO:0009753) and amino (GO:0043200) acids; cellular compartment include cytoplasm (GO:0005737) and cytosol (GO:0005829); and protein binding (GO:0005515 and GO:0042802) as molecular function. Whereas for 30dpi contrast biological process include collagen and cuticulin-based cuticle development (GO:0040002) and innate immune response (GO:0045087); cellular compartment includes nucleus (GO:0005634) and extracellular region (GO:000576); and protein binding (GO:0005515 and GO:0042802) as molecular function.

**Figure 6.**
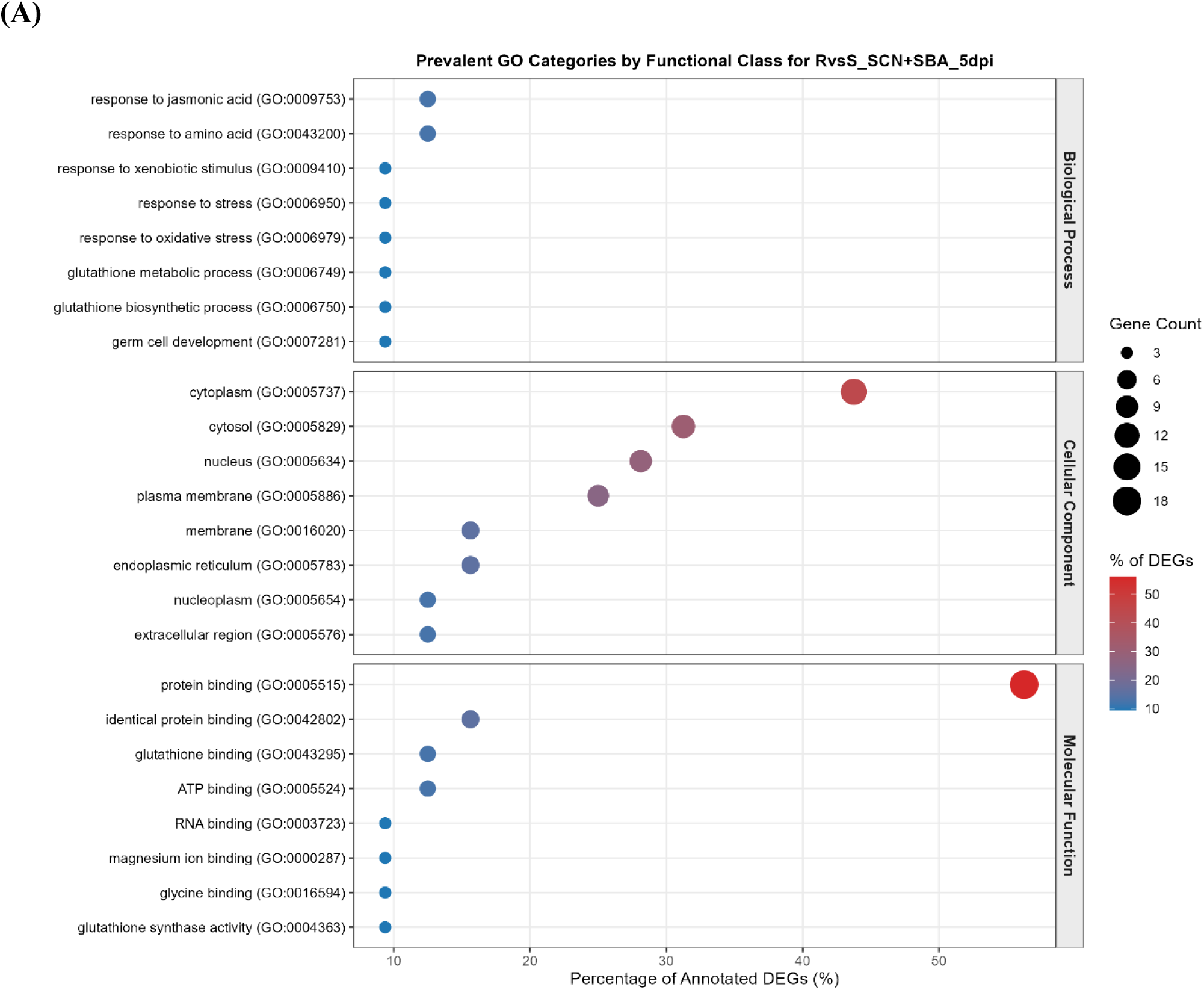
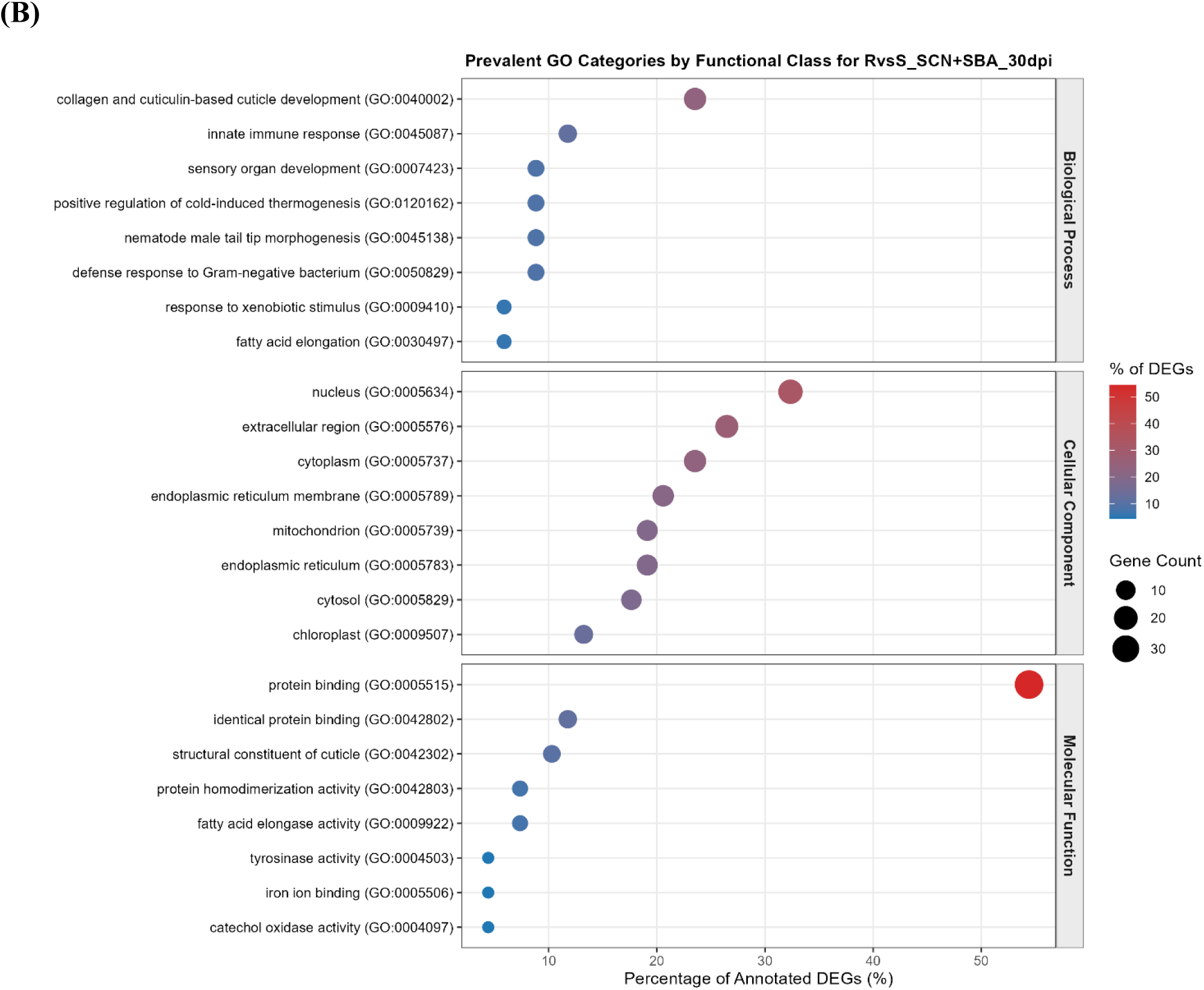
Gene Ontology (GO) annotations for DEGs in the resistant vs susceptible genotype for five and 30 DPI time points. The dotplots illustrates top eight GOs associated with Biological Process, Cellular Component, and Molecular Function for **(A)** SCN + SBA infestation at five and **(B)** SCN + SBA infestation at 30 DPI. The x-axis represents percentage of annotated DEGs, while the y-axis represents the associated GOs. Different color bars represent % of DEGs and circle represents gene counts, as shown in the legend.

The KO enrichment for resistant vs susceptible cultivars treated with SCN + SBA at five DPI flagged K04958 (inositol 1,4,5-triphosphate receptor type-1, n=3) and Hetgly01930.t1 as enriched ortholog and transcript. Similarly, at 30 DPI, K08202 [MFS transporter, OCT family, solute carrier family 22 (organic cation transporter), member 4/5/21; SLC22A4_5_21, OCTN, n= 5), K10203 (elongation of very long chain fatty acids protein 6; ELOVL6, n=5), K11262 (acetyl-CoA carboxylase/biotin carboxylase 1; ACACA, n = 3), and K00505 (tyrosinase; TYR, n=3) were enriched as a KO, and Hetgly12920.t1 (n=9) as a transcript.

## 4. Discussion

We analyzed SCN genes in a three-way interaction between soybean, SCN, and SBA from soybean root transcriptomics, building on previously generated RNA sequencing data by Neupane et. al [12,15,16]. While the majority of studies have focused on parallel transcriptome analysis of individual plant and parasite [45–48]; the decreasing cost of next-generation sequencing and increasing high-depth coverage would essentially capture the parasite’s transcriptomic expression while lodged inside host tissue. To the best of our knowledge, this is the first use of host transcriptomics to study gene expression of parasitic plant nematodes in soybean cultivars.

### 4.1. Asymmetric responsiveness between genotypes

Within-genotype contrasts revealed a striking asymmetry; the susceptible genotype showed essentially no significant transcriptional responses to SCN or SCN + SBA treatment at either timepoint (0 and 2 DEGs across all four contrasts). While the resistant genotype showed a clear, escalating response, culminating in 870 and 1127 DEGs distinguishing SCN- and SCN + SBA-treated roots at five vs 30 DPI. These findings highlight that within the susceptible cultivar, both single and dual pest infestation function equivalently, which is congruent with Neupane et al. [16] results showing an almost unaltered SCN population for treatment contrasts in susceptible soybean. While the same research showed a non-significant SCN population change for the treatment contrasts in the resistant cultivar, the significantly expressed genes reported by this study highlight the potential combative action of SCN trying to overcome soybean resistance.

Additionally, the comparison between susceptible and resistant genotypes showed only three and one significantly expressed genes at five and 30 DPI for SCN treatment. The significantly lower gene expression reported here aligns with previous research highlighting cytological and ultrastructural similarities present in both resistant and susceptible reactions during early response to syncytium [49–51]. However, the SCN + SBA treatment analysis revealed 63 and 105 genes being differentially expressed at five and 30 DPI when comparing between cultivars, respectively. These findings highlight that pathogenicity of SBA potentially weakens the soybean defense response, irrespective of genotype, amplifying SCN pathological action (discussed later). This aligns with McCarville et al. [13], who also reported an increase in SCN population under multiple pathogenic infestations in the resistant soybean cultivar.

The SCN pathogenicity is relevant to the effector molecules (or parasitome) produced in the esophageal glands of nematodes, which are released into the host tissues through the stylet [52]. These molecules contribute to virulence by either overexpressing virulence factor proteins or by repressing ETI (effector-triggered immunity)-inducing proteins [53]. Gao et al. [54] identified 51 SCN parasitome, while later Noon et al. [55] predicted the number to be 82. While the SCN can enter the roots of both susceptible and resistant soybean cultivars [56], resistant cultivars interfere with the SCN life cycle by disrupting syncytium formation, thereby escaping pathogenicity [57]. *Rhg* (resistance to *H. glycines*)-1 and -2 in soybean are the two major QTLs (quantitative trait loci) conferring resistance to SCN [58].

### 4.2. Enrichment analyses

Genes expressed profoundly (n=1,829) were subjected to enrichment analysis to determine their significant functions. Among the three genes upregulated in the resistant vs susceptible comparison when infested with SCN at five DPI K00940 (nucleoside-diphosphate kinase, NDK), K24646 (PR domain zinc finger protein 14, PRDM14), and K08770 (ubiquitin B/C) were enriched. NDK functions as an effector protein affecting host metabolic pathways [59,60] and ubiquitination is potentially involved in host protein degradation [61], whereas PRDM14 ortholog PRDM1 is associated with stress-specific developmental adaptations in *Caenorhabditis elegans* [62]. These findings suggest potential strategies adopted by SCNs for combating the resistant soybean cultivar in early developmental stages. Whereas for SCN + SBA infestation at 5 DPI, K04958 (inositol 1,4,5-triphosphate receptor type-1, ITPR1), K16302 (metal transporter CNNM), and K21456 (glutathione synthase, GSS) were highly enriched in addition to K00940 and K24646 (excluding K08770). ITPR1 and CNNM are involved in Ca^2+^ and Mg^2+^ signaling [63,64], while GSS may promote susceptibility by modulating thiol production, as seen in Arabidopsis infected by *Meloidogyne incognita* [65]. Cotton et al. [66] reported elevated glutathione synthase signaling in the early parasitic stage of *Globodera pallida* infecting potato. Additionally, the report by Jiang et al. [67] during pH- and salt-stressed conditions aligns with our enrichment results. These findings from our study suggest that dual infestation, aided by SBA, potentially promotes SCN activities in resistant cultivars.

At 30 DPI, the contrast between resistant and susceptible cultivars infested with SCN + SBA, with enhanced KO terms: K08202, K10203, K11262, and K00505, signaled membrane transport and defense activities associated with cuticular/suberin barrier reinforcement. The major facilitator superfamily (MFS, K08202), involved in secondary active transport, is the largest known gene superfamily [68]. These genes are involved as solute carrier or organic cation transporters in organisms, including *C. elegans* [69], along with regulating oxidative stress response and development in *Drosophila melanogaster* [70]. While Acetyl-CoA carboxylase (K11262) is involved in fatty acid biosynthesis, catalyzing Acetyl-CoA through ATP-dependent carboxylation [71,72]. Gutbrod et al. [73] reported lipid depletion and growth arrest in nematodes when acetyl-CoA carboxylase is depleted. Additionally, ELOVL6 (K10203) is involved in the fatty acid elongation cycle [74] and the suppression of its ortholog ELO-2 in *C. elegans* affected ultradian rhythms [75]. Tyrosinase (K00505) is one of the nematodes’ effectors modulating plant hormone homeostasis, where Habash et al. [76] showed that gene silencing in *Heterodera schachtii* reduced virulence in Arabidopsis. The canonical enhancement of the genes involved in developmental processes, mainly solute transportation and the cascade of the fatty acid synthesis pathway, during the late infection process suggests potential completion of the nematode life cycle, culminating in egg formation.

Overall, the enrichment analysis highlighted increased SCN activity when combined with SBA. The significantly overexpressed genes in the early infestation stage potentially armed the pest to counteract molecular resistance against resistant soybean cultivars. In contrast, during the late stage, the enriched genes suggested potential dormancy development in the surviving SCN population.

## 5. Conclusion

In this study, we conducted a comprehensive analysis of the transcriptome of SCN utilizing the soybean root RNA sequencing data. We found that 1,829 genes (transcript count > 10) were significantly (padj < 0.05) expressed in the soybean cultivar contrasts during single (SCN only) and combined infestation (SCN + SBA) compared at two time points (five and 30 DPI). While within-genotype contrast for susceptible soybean did not result in significant gene regulation, suggesting similar mode of SCN action for both solo and co-infestation. However, co-infestation in resistant soybean cultivar enhanced the SCN activity with significant differentially expressed genes, enriched in resistance breakdown functions. Overall, the findings serve as baseline data for understanding the molecular pathogenicity of SCN in single and co-infestation with SBA, promoting potential implications in molecular breeding programs to produce soybean genotypes resistant to cyst nematode.

## 6. Limitations

While this study provides baseline data on the impact of SCN on soybean cultivars under single or dual infestation, the findings are based solely on soybean root RNA sequencing to evaluate SCN gene expression at two time points. This potentially limits the visualization of SCN’s early life-stage expression promoting SCN pathogenicity. Moreover, the SCN resistant cultivar used in this study was susceptible to SBA. Therefore, we suggest a rigorous analysis, considering spatial or single-cell or SCN-isolated transcriptomics and time points based on SCN developmental stages, experimenting on various soybean genotypes including SCN-SBA resistant cultivar along with accounting for the profiling of SBA’s gene regulation.

## Acknowledgement

The authors appreciate the help from Emmanuel Byamukama (South Dakota State University), who kindly provided soil infested with *H. glycines* HG Type-0 for the experimentation. We acknowledge the valuable assistance provided by Febina Mathew (North Dakota State University) in experimental design. Additionally, we thank the High Performance Computing (HPC) Cluster at South Dakota State University for providing computational resources.

## Author contributions

B.A. performed the methodological investigation, data analyses, visualization, and drafted, edited, and revised the original manuscript. S.N. conducted the greenhouse experiments, as part of his Ph.D. research, and assisted in editing and reviewing. S.S. contributed in editing and reviewing. M.P.N. and A.J.V. conceived the project, designed the experiments, and revised the manuscript. All authors have read and agreed to the published version of the manuscript.

## Ethics approval and consent to participate

Not applicable.

## Consent for publication

Not applicable.

## Data availability

The genome, cds, and protein sequences of *Heterodera glycines* analyzed in this study are accessible through WormBase Parasite (https://parasite.wormbase.org/Heterodera_glycines_prjna381081/Info/Index). Raw RNA sequencing reads associated with BioProject PRJNA514200 (SRA study/Project ID: SRP178193) are available from the NCBI Sequence Read Archive (SRA). In addition, the processed/transformed transcript abundance matrix, along with the eggnog-mapped orthologs, will be publicly hosted on **Figshare**. The codes and scripts used in this study are publicly available through the authors’ GitHub repository: https://github.com/abibek52/soybean_cyst_nematode_transciptome.

---

**Supplementary Table 1.** Summary statistics of the transcriptomic data used in this study.

| S.N. | Genotype | Treatment | Days | Replication | Number of raw reads | GC % | Trimmed reads | Percentage of clean reads | Mapped reads | Percentage of mapped reads | Accession number |
|---|---|---|---|---|---|---|---|---|---|---|---|
| 1 | PI518671 | Control | 5 | 1 | 17,242,793 | 45 | 17,239,066 | 99.98 | 19,902 | 0.11544709 | SRR8427397 |
| 2 |  |  |  | 2 | 22,062,929 | 46 | 22,055,685 | 99.97 | 118,403 | 0.536836648 | SRR8427395 |
| 3 |  |  |  | 3 | 21,220,300 | 44 | 21,213,623 | 99.97 | 32,485 | 0.15313273 | SRR8427393 |
| 4 |  |  | 30 | 1 | 20,522,473 | 44 | 20,518,044 | 99.98 | 38,546 | 0.187863911 | SRR8427405 |
| 5 |  |  |  | 2 | 28,600,503 | 44 | 28,593,731 | 99.98 | 26,102 | 0.091285744 | SRR8427404 |
| 6 |  |  |  | 3 | 20,577,190 | 44 | 20,570,977 | 99.97 | 21,455 | 0.104297428 | SRR8427407 |
| 7 |  | SCN | 5 | 1 | 20,389,378 | 44 | 20,383,629 | 99.97 | 765,453 | 3.755234164 | SRR8427406 |
| 8 |  |  |  | 2 | 10,518,888 | 44 | 10,516,365 | 99.98 | 382,266 | 3.634963222 | SRR8427401 |
| 9 |  |  |  | 3 | 21,303,947 | 44 | 21,298,111 | 99.97 | 212,118 | 0.995947481 | SRR8427400 |
| 10 |  |  | 30 | 1 | 29,875,777 | 44 | 29,868,305 | 99.97 | 1,317,095 | 4.409674402 | SRR8427366 |
| 11 |  |  |  | 2 | 20,569,129 | 45 | 20,564,513 | 99.98 | 717,133 | 3.487235511 | SRR8427367 |
| 12 |  |  |  | 3 | 23,663,582 | 44 | 23,657,909 | 99.98 | 669,673 | 2.830651686 | SRR8427368 |
| 13 |  | SCN+SBA | 5 | 1 | 21,569,888 | 44 | 21,563,432 | 99.97 | 648,685 | 3.008264176 | SRR8427398 |
| 14 |  |  |  | 2 | 57,520,568 | 44 | 57,503,170 | 99.97 | 5,145,755 | 8.948645788 | SRR8427381 |
| 15 |  |  |  | 3 | 16,889,301 | 45 | 16,883,954 | 99.97 | 745,344 | 4.414510961 | SRR8427382 |
| 16 |  |  | 30 | 1 | 23,727,017 | 45 | 16,882,449 | 71.15 | 170,397 | 1.009314466 | SRR8427364 |
| 17 |  |  |  | 2 | 22,378,982 | 44 | 22,373,777 | 99.98 | 959,069 | 4.2865762 | SRR8427365 |
| 18 |  |  |  | 3 | 27,673,846 | 44 | 27,668,291 | 99.98 | 343,011 | 1.23972601 | SRR8427370 |
| 19 | MN1806CN | Control | 5 | 1 | 18,937,367 | 46 | 18,932,017 | 99.97 | 121,954 | 0.644168025 | SRR8427396 |
| 20 |  |  |  | 2 | 26,710,585 | 43 | 26,702,238 | 99.97 | 118,963 | 0.445516964 | SRR8427394 |
| 21 |  |  |  | 3 | 21,327,385 | 46 | 21,320,799 | 99.97 | 105,686 | 0.495694369 | SRR8427372 |
| 22 |  |  | 30 | 1 | 24,104,763 | 45 | 24,099,789 | 99.98 | 22,515 | 0.093424054 | SRR8427391 |
| 23 |  |  |  | 2 | 32,183,362 | 44 | 32,174,938 | 99.97 | 92,202 | 0.286564655 | SRR8427392 |
| 24 |  | SCN | 5 | 1 | 25,443,012 | 44 | 25,435,147 | 99.97 | 845,300 | 3.323354097 | SRR8427379 |
| 25 |  |  |  | 2 | 20,043,049 | 45 | 20,037,212 | 99.97 | 607,619 | 3.032452818 | SRR8427380 |
| 26 |  |  |  | 3 | 9,847,269 | 45 | 9,844,767 | 99.97 | 652,135 | 6.62417912 | SRR8427377 |
| 27 |  |  | 30 | 1 | 25,200,882 | 43 | 25,192,664 | 99.97 | 624,843 | 2.480257745 | SRR8427371 |
| 28 |  |  |  | 2 | 22,192,100 | 43 | 22,186,459 | 99.97 | 157,013 | 0.707697429 | SRR8427383 |
| 29 |  |  |  | 3 | 20,653,286 | 43 | 20,648,111 | 99.97 | 133,315 | 0.645652283 | SRR8427384 |
| 30 |  | SCN+SBA | 5 | 1 | 20,636,498 | 44 | 20,630,026 | 99.97 | 179,865 | 0.871860268 | SRR8427373 |
| 31 |  |  |  | 2 | 22,488,050 | 44 | 22,482,625 | 99.98 | 434,499 | 1.932599063 | SRR8427374 |
| 32 |  |  |  | 3 | 22,033,213 | 45 | 22,028,303 | 99.98 | 411,780 | 1.869322389 | SRR8427408 |
| 33 |  |  | 30 | 1 | 19,498,275 | 43 | 19,491,491 | 99.97 | 114,297 | 0.58639434 | SRR8427388 |
| 34 |  |  |  | 2 | 27,765,044 | 44 | 27,759,095 | 99.98 | 263,425 | 0.948968257 | SRR8427389 |
| 35 |  |  |  | 3 | 43,325,617 | 44 | 43,312,161 | 99.97 | 259,666 | 0.599522153 | SRR8427390 |

## References

1. Medic J, Atkinson C, Hurburgh CR (2014). Current knowledge in soybean composition. Journal of the American Oil Chemists’ Society 91:363–384. 10.1007/s11746-013-2407-9

2. Bent AF (2022). Exploring soybean resistance to soybean cyst nematode. Annual Review of Phytopathology 60:379–409. 10.1146/annurev-phyto-020620-120823

3. USDA-FAS (2026). United States Department of Agriculture Foreign Agricultural Service. Productions - soybeans. (Last accessed on August 26, 2026). Available at: https://www.fas.usda.gov/data/production/2222000.

4. Allen T, Mueller D, Sisson A, Sikora E, Faske T, Betts A, Dufault N, Grabau Z, Kemerait B, Camiletti B, Telenko, Onofre R, Bradley C, Padgett B, Price T, Watson T, Chilvers M, Malvick D, Bish M, Mangel D, Bergstrom G, Lux L, Mathew F, Webster W, Lopez-Nicora H, Duffeck M, Tenuta A, Collins A, Esker P, Roth G, Mueller J, Plumblee M, Shires M, Kelly H, Isakeit T, Langston D, Zeng Y, Smith D (2025). Soybean disease loss estimates from the United States and Ontario, Canada—2025. Crop Protection Network CPN-1018-25: 10.31274/cpn-20260306-0

5. Wang J, Niblack TL, Tremain JA, Wiebold WJ, Tylka GL, Marett CC, Noel GR, Myers O, Schmidt ME (2003). Soybean cyst nematode reduces soybean yield without causing obvious aboveground symptoms. Plant Disease 87:623–628. 10.1094/PDIS.2003.87.6.623

6. Hartman GL, Domier LL, Wax LM, Helm CG, Onstad DW, Shaw JT, Solter LF, Voegtlin DJ, D’Arcy CJ, Gray ME, Steffey KL, Isard SA, Orwick PL (2001). Occurrence and distribution of *Aphis glycines* on soybeans in Illinois in 2000 and its potential control. Plant Health Progress 2:17. 10.1094/PHP-2001-0205-01-HN

7. Telenko DEP, Allen TW, Sisson AJ, Bergstrom GC, Betts AK, Bish MD, Bissonnette KM, Bond JP, Bonkowski J, Bradley CA, Byamukama E, Camiletti BX, Chilvers MI, Collins AA, Damicone JP, Dufault NS, Duffeck MR, Esker PD, Faske TR, Grabau ZJ, Harbach CJ, Isakeit T, Jackson-Ziems TA, Kelly HM, Kemerait RC, Kleczewski NM, Langston DB, Lofton J, Lopez-Nicora HD, Lux LR, Malvick DK, Mangel DJ, Markell SG, Mathew FM, Mehl HL, Mehl KM, Mideros SX, Mueller DS, Mueller JD, Nelson BD, Onofre RB, Padgett GB, Plumblee MT, Price PP, Shires MK, Sikora EJ, Small I, Smith DL, Spurlock TN, Tande CA, Tenuta AU, Thiessen LD, Warner F, Watson TT, Webster RW, Wise KL, Zeng Y (2025). Estimated soybean yield and economic losses caused by diseases in the United States and Ontario, Canada, from 2020 to 2024. Plant Health Progress 27:269–282. 10.1094/PHP-09-25-0227-RS

8. Tylka G (2023). New data-based calculator estimates economic yield loss from SCN in individual fields. Integrated Crop Management. Iowa State University Extension and Outreach. (Last accessed on September 9, 2026). Available at: https://crops.extension.iastate.edu/cropnews/2023/11/new-data-based-calculator-estimates-economic-yield-loss-scn-individual-fields.

9. Kim CS, Schaible G, Garrett L, Lubowski R, Lee D (2008). Economic impacts of the U. S. soybean aphid infestation: A multi-regional competitive dynamic analysis. Agricultural and Resource Economics Review 37:227–242. 10.1017/S1068280500003026

10. Natukunda MI, MacIntosh GC (2020). The resistant soybean-*Aphis glycines* interaction: Current knowledge and prospects. Frontiers in Plant Science Volume 11–2020: 10.3389/fpls.2020.01223

11. Tylka G, Gebhart GD, Marett CC, Mullaney MP, Rasmussen JT (2025). Evaluation of soybean varieties resistant to soybean cyst nematode in Iowa, 2025. 10.31274/usp-20260507-3

12. Neupane S, Purintun JM, Mathew FM, Varenhorst AJ, Nepal MP (2019). Molecular basis of soybean resistance to soybean aphids and soybean cyst nematodes. Plants 8:374. 10.3390/plants8100374

13. McCarville MT, O’Neal M, Tylka GL, Kanobe C, MacIntosh GC (2012). A nematode, fungus, and aphid interact via a shared host plant: implications for soybean management. Entomologia Experimentalis et Applicata 143:55–66. 10.1111/j.1570-7458.2012.01227.x

14. McCarville MT, Soh DH, Tylka GL, O’Neal ME (2014). Aboveground feeding by soybean aphid, *Aphis glycines*, affects soybean cyst nematode, *Heterodera glycines*, reproduction belowground. PLOS ONE 9:e86415. 10.1371/journal.pone.0086415

15. Neupane S, Mathew FM, Varenhorst AJ, Nepal MP (2019). Transcriptome profiling of interaction effects of soybean cyst nematodes and soybean aphids on soybean. Scientific Data 6:133. 10.1038/s41597-019-0140-4

16. Neupane S, Varenhorst AJ, Nepal MP (2026). Time- and genotype-dependent root-transcriptomic responses of soybean to combined soybean aphid and soybean cyst nematode infestation. Plants 15:2014. 10.3390/plants15132014

17. Andrews S (2023). FastQC: A quality control tool for high throughput sequence data. Available at: https://www.bioinformatics.babraham.ac.uk/projects/fastqc/.

18. Ewels P, Magnusson M, Lundin S, Käller M (2016). MultiQC: summarize analysis results for multiple tools and samples in a single report. Bioinformatics 32:3047–3048. 10.1093/bioinformatics/btw354

19. Kong Y (2011). Btrim: A fast, lightweight adapter and quality trimming program for next-generation sequencing technologies. Genomics 98:152–153. 10.1016/j.ygeno.2011.05.009

20. Sternberg PW, Van Auken K, Wang Q, Wright A, Yook K, Zarowiecki M, Arnaboldi V, Becerra A, Brown S, Cain S, Chan J, Chen WJ, Cho J, Davis P, Diamantakis S, Dyer S, Grigoriadis D, Grove CA, Harris T, Howe K, Kishore R, Lee R, Longden I, Luypaert M, Müller H-M, Nuin P, Quinton-Tulloch M, Raciti D, Schedl T, Schindelman G, Stein L (2024). WormBase 2024: status and transitioning to Alliance infrastructure. Genetics 227:iyae050. 10.1093/genetics/iyae050

21. Patro R, Duggal G, Love MI, Irizarry RA, Kingsford C (2017). Salmon provides fast and bias-aware quantification of transcript expression. Nat Methods 14:417–419. 10.1038/nmeth.4197

22. Grüning B, Dale R, Sjödin A, Chapman BA, Rowe J, Tomkins-Tinch CH, Valieris R, Köster J, The Bioconda T (2018). Bioconda: sustainable and comprehensive software distribution for the life sciences. Nature Methods 15:475–476. 10.1038/s41592-018-0046-7

23. Pagès H, Carlson M, Aboyoun P, Falcon S, Morgan M (2026). txdbmaker: Tools for making TxDb objects from genomic annotations. In. Available at: https://bioconductor.org/packages/txdbmaker. 10.18129/B9.bioc.txdbmaker

24. Pagès H, Carlson M, Falcon S, Li N (2026). AnnotationDbi: Manipulation of SQLite-based annotations in Bioconductor. R package version 1.74.0. Available at: https://bioconductor.org/packages/AnnotationDbi. 10.18129/B9.bioc.AnnotationDbi.

25. Huber W, Carey VJ, Gentleman R, Anders S, Carlson M, Carvalho BS, Bravo HC, Davis S, Gatto L, Girke T, Gottardo R, Hahne F, Hansen KD, Irizarry RA, Lawrence M, Love MI, MacDonald J, Obenchain V, Oleś AK, Pagès H, Reyes A, Shannon P, Smyth GK, Tenenbaum D, Waldron L, Morgan M (2015). Orchestrating high-throughput genomic analysis with Bioconductor. Nature Methods 12:115–121. 10.1038/nmeth.3252

26. R_Core_Team (2026). R: A language and environment for statistical computing. R Foundation for Statistical Computing, Vienna, Austria 10.32614/R.manuals

27. Benjamini Y, Hochberg Y (1995). Controlling the false discovery rate: A practical and powerful approach to multiple testing. Journal of the Royal Statistical Society: Series B (Methodological) 57:289–300. 10.1111/j.2517-6161.1995.tb02031.x

28. Soneson C, Love MI, Robinson MD (2015). Differential analyses for RNA-seq: transcript-level estimates improve gene-level inferences. F1000Res 4:1521. 10.12688/f1000research.7563.2

29. Love MI, Huber W, Anders S (2014). Moderated estimation of fold change and dispersion for RNA-seq data with DESeq2. Genome Biology 15:550. 10.1186/s13059-014-0550-8

30. Kolde R (2015). Pheatmap: Pretty heatmaps. R package version 1.0.13. Available at: https://pheatmap.com/. 10.32614/CRAN.package.pheatmap

31. Wickham H (2016). ggplot2: Elegant graphics for data analysis, 2 edn. Use R! Springer Cham. 10.1007/978-3-319-24277-4

32. Auguie B (2026). gridExtra: Miscellaneous functions for “Grid” graphics. R package version 2.3.1. 10.32614/CRAN.package.gridExtra

33. Hadley W, Davis V, Maximilian G (2025). tidyr: Tidy messy data. R package version 1.3.2. Available at: https://CRAN.R-project.org/package=tidyr. 10.32614/CRAN.package.tidyr

34. Hadley W, Romain F, Lionel H, Kirill M, Davis V (2026). dplyr: A grammar of data manipulation. R package version 1.2.1. Available at: https://CRAN.R-project.org/package=dplyr. 10.32614/CRAN.package.dplyr

35. Neuwirth E (2022). RColorBrewer: Colorbrewer palettes. R package version 1.1–3. 10.32614/CRAN.package.RColorBrewer

36. Chen H, Boutros PC (2011). VennDiagram: a package for the generation of highly-customizable Venn and Euler diagrams in R. BMC Bioinformatics 12:35. 10.1186/1471-2105-12-35

37. Hervé P, Patrick A, Robert G, Saikat D (2026). Biostrings: Efficient manipulation of biological strings. R package version 2.80.1. Available at: https://bioconductor.org/packages/Biostrings. 10.18129/B9.bioc.Biostrings

38. Cantalapiedra CP, Hernández-Plaza A, Letunic I, Bork P, Huerta-Cepas J (2021). eggNOG-mapper v2: Functional annotation, orthology assignments, and domain prediction at the metagenomic scale. Mol Biol Evol 38:5825–5829. 10.1093/molbev/msab293

39. Hernández-Plaza A, Deng Z, Robledo-Yagüe F, Szklarczyk D, von Mering C, Bork P, Huerta-Cepas J (2026). eggNOG v7: Phylogeny-based orthology predictions and functional annotations. Nucleic Acids Research 54:D402–D408. 10.1093/nar/gkaf1249

40. Buchfink B, Reuter K, Drost H-G (2021). Sensitive protein alignments at tree-of-life scale using DIAMOND. Nature Methods 18:366–368. 10.1038/s41592-021-01101-x

41. Hadley W, Bryan J (2026). readxl: Read excel files. R package version 1.6.0. Available at: https://readxl.tidyverse.org/

42. Hadley W (2025). stringr: Simple, consistent wrappers for common string operations. R package version 1.6.0. Available at: https://stringr.tidyverse.org/

43. Carlson M (2026). GO.db: A set of annotation maps describing the entire Gene Ontology. R package version 3.23.1. Available at: https://bioconductor.org/packages/GO.db. 10.18129/B9.bioc.GO.db

44. Kanehisa M, Furumichi M, Sato Y, Matsuura Y, Ishiguro-Watanabe M (2025). KEGG: biological systems database as a model of the real world. Nucleic Acids Research 53:D672–D677. 10.1093/nar/gkae909

45. Ithal N, Recknor J, Nettleton D, Hearne L, Maier T, Baum TJ, Mitchum MG (2007). Parallel genome-wide expression profiling of host and pathogen during soybean cyst nematode infection of soybean. Molecular Plant-Microbe Interactions 20:293–305. 10.1094/MPMI-20-3-0293

46. Wan J, Vuong T, Jiao Y, Joshi T, Zhang H, Xu D, Nguyen HT (2015). Whole-genome gene expression profiling revealed genes and pathways potentially involved in regulating interactions of soybean with cyst nematode (*Heterodera glycines* Ichinohe). BMC Genomics 16:148. 10.1186/s12864-015-1316-8

47. Ste-Croix DT, St-Marseille A-FG, Lord E, Bélanger RR, Brodeur J, Mimee B (2020). Genomic profiling of virulence in the soybean cyst nematode using single-nematode sequencing. Phytopathology® 111:137–148. 10.1094/PHYTO-08-20-0348-FI

48. Bai P-P, Jia X-Y, Chen L, Han Y, Han S, Weng L, Feng X (2025). Single-nucleus transcriptome analyses uncover a dynamic transcriptional landscape of soybean roots in response to soybean cyst nematode infection. bioRxiv 2025.2008.2006.668855. 10.1101/2025.08.06.668855

49. Klink VP, Martins VE, Alkharouf NW, Overall CC, MacDonald MH, Matthews BF (2007). A decline in transcript abundance for *Heterodera glycines* homologs of *Caenorhabditis elegans* uncoordinated genes accompanies its sedentary parasitic phase. BMC Developmental Biology 7:35. 10.1186/1471-213X-7-35

50. Klink VP, Overall CC, Alkharouf NW, MacDonald MH, Matthews BF (2007). Laser capture microdissection (LCM) and comparative microarray expression analysis of syncytial cells isolated from incompatible and compatible soybean (*Glycine max*) roots infected by the soybean cyst nematode (*Heterodera glycines*). Planta 226:1389–1409. 10.1007/s00425-007-0578-z

51. Klink VP, Matthews BF (2009). Emerging approaches to broaden resistance of soybean to soybean cyst nematode as supported by gene expression studies. Plant Physiology 151:1017–1022. 10.1104/pp.109.144006

52. Mitchum MG, Hussey RS, Baum TJ, Wang X, Elling AA, Wubben M, Davis EL (2013). Nematode effector proteins: an emerging paradigm of parasitism. New Phytologist 199:879–894. 10.1111/nph.12323

53. Sonah H, Zhang X, Deshmukh RK, Borhan MH, Fernando WGD, Bélanger RR (2016). Comparative transcriptomic analysis of virulence factors in *Leptosphaeria maculans* during compatible and incompatible interactions with canola. Frontiers in Plant Science Volume 7–2016: 10.3389/fpls.2016.01784

54. Gao B, Allen R, Maier T, Davis EL, Baum TJ, Hussey RS (2003). The parasitome of the phytonematode *Heterodera glycines*. Molecular Plant-Microbe Interactions 16:720–726. 10.1094/MPMI.2003.16.8.720

55. Noon JB, Hewezi T, Maier TR, Simmons C, Wei J-Z, Wu G, Llaca V, Deschamps S, Davis EL, Mitchum MG, Hussey RS, Baum TJ (2015). Eighteen new candidate effectors of the Phytonematode *Heterodera glycines* produced specifically in the secretory esophageal gland cells during parasitism. Phytopathology® 105:1362–1372. 10.1094/PHYTO-02-15-0049-R

56. Verma A, Lee C, Morriss S, Odu F, Kenning C, Rizzo N, Spollen WG, Lin M, McRae AG, Givan SA, Hewezi T, Hussey R, Davis EL, Baum TJ, Mitchum MG (2018). The novel cyst nematode effector protein 30D08 targets host nuclear functions to alter gene expression in feeding sites. New Phytologist 219:697–713. 10.1111/nph.15179

57. Sobczak M, Avrova A, Jupowicz J, Phillips MS, Ernst K, Kumar A (2005). Characterization of susceptibility and resistance responses to potato cyst nematode (*Globodera* spp.) infection of tomato lines in the absence and presence of the broad-spectrum nematode resistance hero gene. Molecular Plant-Microbe Interactions 18:158–168. 10.1094/MPMI-18-0158

58. Mitchum MG (2016). Soybean resistance to the soybean cyst nematode *Heterodera glycines*: An update. Phytopathology® 106:1444–1450. 10.1094/PHYTO-06-16-0227-RVW

59. Wang Z, Ge J-Q, Chen H, Cheng X, Yang Y, Li J, Whitworth RJ, Chen M-S (2018). An insect nucleoside diphosphate kinase (NDK) functions as an effector protein in wheat - Hessian fly interactions. Insect Biochemistry and Molecular Biology 100:30–38. 10.1016/j.ibmb.2018.06.003

60. Gounaris K (2002). Nucleotidase cascades are catalyzed by secreted proteins of the parasitic nematode *Trichinella spiralis*. Infection and Immunity 70:4917–4924. 10.1128/iai.70.9.4917-4924.2002

61. Vasquez YM, Romero MF, Bowers RM, Rohwer RR, McMahon KD, Woyke T, Schulz F (2026). Vicennial metagenomic time series unveils evolutionary dynamics of giant viruses in a freshwater ecosystem. Nature Communications 17:7192. 10.1038/s41467-026-73437-x

62. Hyun M, Kim J, Dumur C, Schroeder FC, You Y-J (2016). BLIMP-1/BLMP-1 and metastasis-associated protein regulate stress resistant development in *Caenorhabditis elegans*. Genetics 203:1721–1732. 10.1534/genetics.116.190793

63. Allman E, Thyagarajan B, Nehrke K (2012). The inositol 1,4,5-trisphosphate receptor in *C. elegans*. Wiley Interdisciplinary Reviews: Membrane Transport and Signaling 1:321–328. 10.1002/wmts.25

64. Ishii T, Funato Y, Hashizume O, Yamazaki D, Hirata Y, Nishiwaki K, Kono N, Arai H, Miki H (2016). Mg2+ extrusion from intestinal epithelia by CNNM proteins Is essential for gonadogenesis via AMPK-TORC1 signaling in *Caenorhabditis elegans*. PLOS Genetics 12:e1006276. 10.1371/journal.pgen.1006276

65. Hasan MS, Damm A, Ijaz MU, Chopra D, Koprivova A, Kopriva S, Grundler FMW, Siddique S (2025). Glutathione promotes susceptibility to *Meloidogyne incognita* by modulating thiol homeostasis in *Arabidopsis*. Phytopathology® 116:320–326. 10.1094/PHYTO-03-25-0090-R

66. Cotton JA, Lilley CJ, Jones LM, Kikuchi T, Reid AJ, Thorpe P, Tsai IJ, Beasley H, Blok V, Cock PJA, den Akker SE-v, Holroyd N, Hunt M, Mantelin S, Naghra H, Pain A, Palomares-Rius JE, Zarowiecki M, Berriman M, Jones JT, Urwin PE (2014). The genome and life-stage specific transcriptomes of *Globodera pallida* elucidate key aspects of plant parasitism by a cyst nematode. Genome Biology 15:R43. 10.1186/gb-2014-15-3-r43

67. Jiang Y, Huang M, Qin R, Jiang D, Chang D, Xie Y, Li C, Wang C (2023). Full-length transcriptome analysis of soybean cyst nematode (*Heterodera glycines*) reveals an association of behaviors in response to attractive pH and salt solutions with activation of transmembrane receptors, ion channels, and Ca2+ transporters. Journal of Agricultural and Food Chemistry 71:8778–8796. 10.1021/acs.jafc.3c00908

68. Drew D, North RA, Nagarathinam K, Tanabe M (2021). Structures and general transport mechanisms by the major facilitator superfamily (MFS). Chemical Reviews 121:5289–5335. 10.1021/acs.chemrev.0c00983

69. Wu X, Fei Y-J, Huang W, Chancy C, Leibach FH, Ganapathy V (1999). Identity of the *F52F12.1* gene product in *Caenorhabditis elegans* as an organic cation transporter. Biochimica et Biophysica Acta (BBA) - Biomembranes 1418:239–244. 10.1016/S0005-2736(99)00020-6

70. Engelhart DC, Azad P, Ali S, Granados JC, Haddad GG, Nigam SK (2020). *Drosophila* SLC22 orthologs related to OATs, OCTs, and OCTNs regulate development and responsiveness to oxidative stress. International Journal of Molecular Sciences 21:2002. 10.3390/ijms21062002

71. Wakil SJ, Titchener EB, Gibson DM (1958). Evidence for the participation of biotin in the enzymic synthesis of fatty acids. Biochimica et Biophysica Acta 29:225–226. 10.1016/0006-3002(58)90177-X

72. Tong L (2013). Structure and function of biotin-dependent carboxylases. Cellular and Molecular Life Sciences 70:863–891. 10.1007/s00018-012-1096-0

73. Gutbrod P, Gutbrod K, Nauen R, Elashry A, Siddique S, Benting J, Dörmann P, Grundler FMW (2020). Inhibition of acetyl-CoA carboxylase by spirotetramat causes growth arrest and lipid depletion in nematodes. Scientific Reports 10:12710. 10.1038/s41598-020-69624-5

74. Naganuma T, Kihara A (2014). Two modes of regulation of the fatty acid elongase ELOVL6 by the 3-ketoacyl-CoA reductase KAR in the fatty acid elongation cycle. PLOS ONE 9:e101823. 10.1371/journal.pone.0101823

75. Kniazeva M, Sieber M, McCauley S, Zhang K, Watts JL, Han M (2003). Suppression of the ELO-2 FA elongation activity results in alterations of the fatty acid composition and multiple physiological defects, including abnormal ultradian rhythms, in *Caenorhabditis elegans*. Genetics 163:159–169. 10.1093/genetics/163.1.159

76. Habash SS, Radakovic ZS, Vankova R, Siddique S, Dobrev P, Gleason C, Grundler FMW, Elashry A (2017). *Heterodera schachtii* tyrosinase-like protein - a novel nematode effector modulating plant hormone homeostasis. Scientific Reports 7:6874. 10.1038/s41598-017-07269-7

